# Echocardiographic evaluation of normal adult left Ventricular geometry in a Nigerian population

**DOI:** 10.1101/2020.04.09.033993

**Authors:** Daniel Chimuanya Ugwuanyi, Joseph Chukwuemeka Eze, Hyacienth Uche Chiegwu, Charles Ugwoke Eze, Chukwudi Thaddeus Nwagbara

**Author notes:** CORRESPONDENT AUTHOR: D.C Ugwuanyi. These authors contributed equally to this work (Joseph Chukwuemeka Eze, Hyacienth Uche Chiegwu). These authors also contributed equally to this work (Charles Ugwoke Eze, Chukwudi Thaddeus Nwagbara).

## Abstract

**Background:** Differences have shown to exist in some echocardiographic measurements that were attributed to racial, ethnic and gender. This study determined echocardiographic baseline data of normal adult left ventricular (LV) geometry in our locality.

**Methods:** The study was performed on 1,192 apparently healthy adults. Participants below the age of 18 years or those with congenital or acquired cardiac abnormalities and history of long-term regular physical training were excluded. Trans-thoracic echocardiography was performed with Vivid T8 GE dedicated echocardiography machine with probe frequency of 1.7 to 3.2 MHz with integrated electrocardiography (ECG) recording electrodes. The study determined normal dimensions of interventricular diamensions. All measurements were indexed to body surface area (BSA) to obtain echocardiographic baseline normal reference values.

**Results:** The mean ± SD values of LV parameters for male and female participants were: LVIDd (44.80 ± 5.71 mm vs 42.75 ± 5.21 mm), LVIDs (33.54 ± 5.37 mm vs 30.38 ± 4.81 mm), and LVPWd (8.32 ± 1.26 mm vs 7.51 ± 1.22 mm). Females had more statistically significant interventricular septum in diastole (IVSd) (8.20 ± 1.38 mm vs 7.05 ± 1.27 mm) and interventricular septum in systole (IVSs) (9.08 ± 1.42 mm vs 8.99 ± 1.33 mm) (P < 0.05).

**Conclusion:** This research established echocardiographic baseline normal adult left ventricular geometry in the study population because in order to detect abnormalities, accurate definition of normal values of echocardiographic measurements is of utmost importance for a reliable clinical decision making.

## 1. Introduction

Measurement of the size of the left ventricle of the heart provides diagnostic clues and prognostic information, and enables the clinician to follow patients in respect of disease progression or improvement [1]. Echocardiography is the most widely used non-invasive imaging tool for the assessment of the heart structure and functions [2]. Various measurements can be performed to determine the size, diameter, length and area of the left ventricle. These measurements are performed at distinct time points of the cardiac cycle, i.e the end of diastole and the end of systole. The end of diastole is seen when the volume of the ventricle is largest, shortly before the mitral valve closes and the mitral annulus descends [3, 4]. The end of systole is the time when the volume of the ventricle is smallest, shortly before the mitral valve opens. Therefore, in order to detect abnormalities, accurate definition of normal values of echocardiographic measurements is of utmost importance in order to be a reliable guide for decision making [3, 4]. Therefore the aim of this study is to determine echocardiographic baseline data of normal adult left ventricular (LV) geometry in Nnewi community setting of Anambra state, Nigeria.

Every effort has been made in establishing normal cardiac dimensions. Such diagnoses include macroscopic examination, genetic studies, electrocardiography, cardiac magnetic resonance imaging (MRI), histological studies and echocardiography [5].

Echocardiography remains the gold standard to determine the structure and fuction of the heart [2]. It is also readily available and cheap when compared to other imaging modalities in Nigeria. The normal baseline values of echocardiographic measured left ventricule of the heart in the literature are mostly based on Caucasian populations [6]. Moreover, there is paucity of literature on the normal echocardiographic values of the left ventricle among adults in our locality. It is of great clinical importance to differentiate between normal and abnormal left ventricles and to explore the normal anatomical variation of these parameters.

This will prove more difficult if the ranges of normal values are unknown for each clime. Therefore, the aim of this study is to evaluate specific normal reference values of adult left ventricular geometry in our locality in a cross sectional study. The findings will be useful in understanding the pathological changes, treatment and management of cardiac patients in the locality.

## 2. Materials and methods

A prospective cross-sectional design was adopted for this study. It was carried out on 1,192 apparently healthy volunteers above 18 years of age who presented for echocardiography study at our center between March 2017 and August 2018. Participants were excluded if they had congenital or acquired cardiac abnormality, had history of long-term regular physical training and If they had any systemic disease (endocrine, collagen, metabolic, nutritional or infectious), hypertension, diabetes mellitus, chronic kidney diseases, and/or chest disease. The participants who wished their relatives to witness the procedure were obliged. This is to give the participants confidence so as to achieve maximum co-operation for better quality results.

Weights were measured with commercially available weighing scale (Hana model). Heights were measured by a meter rule. Height and weight were used to estimate body mass index (BMI) in kg/m^2^ and body surface area (BSA) in m^2^. Where BSA = square root of ([height in cm x weight in kg]/3600).

**Sample size estimation**

The minimum sample size for this study is given by:

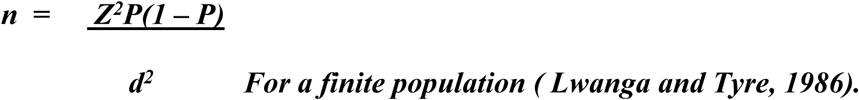

Where:

n = minimum Sample size.

Z = 1.96 at 95% confidence interval, that is the standardized Z – score.

P = estimated population proportion.

Since this proportion for the population under study is not known, a value of 50% (0.5) is assigned to obtain maximum value for P.

d = absolute precision required on either side of the proportion = 50% (0.05).

Substituting the values of z, p and d in the above equation:

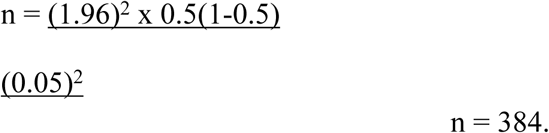

Therefore in order to minimize sample error, sample size of 1,192 apparently healthy subjects was used for the study.

**Scanning protocol or technique:** The participants were scanned using 2D, M-mode and Doppler measurements. Standard trans-thoracic echocardiographic studies with machine- integrated ECG recording were performed using Vivid T8 GE machine with sector probe of frequency range from 1.7 to 3.2 MHz. The choice of the probe was to get adequate visualization of the heart through the intercoastal space. All studies were done with patients lying in the left lateral decubitus position and breathing quietly [7, 8]. Ultrasound gel was applied to ensure proper coupling of the transducer and good transmission of the ultrasound beam into the subjects. From the parasternal window, parasternal long axis views were obtained by placing the transducer in the left third or fourth intercostal space adjacent to the sternum with the knob pointing toward the right shoulder. After confirming a true long axis view that was perpendicular to the centre of the true long axis of the left ventricle (LV), M-mode image was taken between the papillary muscle and at the tip of the mitral valve [9]. Measurements were made from the leading edge of the septal endocardium to the leading edge of posterior wall endocardium [8]. Measurements for the interventricular septum at end diastole (IVSd), LV internal dimensions at end diastole (LVEDd) and LV posterior wall thickness at end diastole (LVPWd) were obtained. Also the measurements were obtained at end systole for each of the parameters mentioned above. All measurements were done on a frozen M-Mode image.

All measurements were divided by BSA to obtain indexed measurements [2, 6].

## 3. Statistical analysis

The data collected were analyzed using Microsoft’s™) Statistical Package for Social Sciences (SPSS) version 20. Descriptive statistics such as mean, standard deviation, and frequencies were generated and calculated for Interventricular septum, left ventricular internal diameter, and left ventricular posterior wall for all age groups. The level of statistical significance was at (*P* < 0.05). Pearson’s correlation and regression analysis were used to determine the relationship between left ventricular parameters with age, height, body weight, body mass index, and body surface area. The relationship between left ventricular baseline values with gender was also determined using correlation and regression analyses. The t-test for two samples assuming equal variance analysis was carried out to assess gender differences in left ventricular parameters. The values of left ventricular parameters generated in the study was compared with the previous studies in Caucasian population to assess inter racial differences using t-test for equality of mean.

## 4. Results and discussion

### 4.1 Social demographic variables

**Table 1**. Presents the age and sex of the participants. The mean age was 57.44 ± 19.10 years. 53.6% of these subjects were males while 46.4% were females.

**Table 1:** Age and Sex distribution of the participants (n = 1,192)

| | F(%) | mean age ± SD<br>$57.44 \pm 19.10$ | | |
| --- | --- | --- | --- | --- |
| Age (years) | frequency |  |  |  |
| 18-22 | 49 |  |  |  |
| 23-27 | 54 |  |  |  |
| 28-32 | 64 |  |  |  |
| 33-37 | 51 |  |  |  |
| 38-42 | 66 |  |  |  |
| 43-47 | 75 |  |  |  |
| 48-52 | 82 |  |  |  |
| 53-57 | 98 |  |  |  |
| 58-62 | 143 |  |  |  |
| 63-67 | 113 |  |  |  |
| 68-72 | 136 |  |  |  |
| 73-77 | 74 |  |  |  |
| 78-82 | 63 |  |  |  |
| 83-87 | 61 |  |  |  |
| 88-93 | 63 |  |  |  |
| Sex |  |  |  |  |
| Male | 639(53.6) |  |  |  |
| Female | 553(46.4) |  |  |  |

**Table 2**. Presents the height and weight of the participants. The mean height was 1.69 ± 0.12m.The mean weight was 75.95 ± 15.74kg.

**Table 2:** Height and Weight of the participants (n = 1,192).

|  | Normal Subjects (n = 1192) |  |  |  |
| --- | --- | --- | --- | --- |
| Range | F(%) | mean height ±<br>SD = $1.69 \pm 0.12\text{m}$ | | |
| **Height (m)** |  |  |  |  |
| **Range** |  |  |  |  |
| 1.16-1.20 | - |  |  |  |
| 1.35-1.40 | 5(0.4) |  |  |  |
| 1.41-1.45 | 11(0.9) |  |  |  |
| 1.46-1.50 | 65(5.5) |  |  |  |
| 1.51-1.55 | 88(7.4) |  |  |  |
| 1.56-1.60 | 97(8.1) |  |  |  |
| 1.61-1.65 | 184(15.4) |  |  |  |
| 1.66-1.70 | 185(15.5) |  |  |  |
| 1.71-1.75 | 172(14.4) |  |  |  |
| 1.76-1.80 | 170(14.3) |  |  |  |
| 1.81-1.85 | 104(8.7) |  |  |  |
| 1.86-1.90 | 59(4.9) |  |  |  |
| 1.91-1.95 | 52(4.4) |  |  |  |
| **Weight (kg)** |  |  |  |  |
| **Range** | F % | mean weight ±<br>SD 75.95<br>$\pm 15.74\text{ kg}$ | | |
| 41-45 | 1(0.1) |  |  |  |
| 46-50 | 62(5.2) |  |  |  |
| 51-55 | 67(5.6) |  |  |  |
| 56-60 | 96(8.1) |  |  |  |
| 61-65 | 103(8.6) |  |  |  |
| 66-70 | 132(11.1) |  |  |  |
| 71-75 | 161(13.5) |  |  |  |
| 76-80 | 135(11.3) |  |  |  |
| 81-85 | 99(8.3) |  |  |  |
| 86-90 | 87(7.3) |  |  |  |
| 91-95 | 87(7.3) |  |  |  |
| 96-100 | 68(5.7) |  |  |  |
| 101-105 | 51(4.3) |  |  |  |
| 106-110 | 43(3.6) |  |  |  |

**Table 3**. Presents the BMI and BSA of the participants. The mean BMI was 26.48 ± 5.15 kg/m^2^. The mean BSA was 1.88 ± 0.24 m^2^.

**Table 3:** BMI and BSA of the participants (n = 1,192).

|  | Normal subjects (n = 1192) |  |  |  |
| --- | --- | --- | --- | --- |
| | F(%) | mean BMI ±<br>SD 26.48 ±<br>5.15 kg/m<sup>2</sup> | | |
| **BMI (Kg/m<sup>2</sup>)** |  |  |  |  |
| Underweight | 11 (0.9) |  |  |  |
| Normal weight | 530 (44.5) |  |  |  |
| Overweight | 429 (36.0) |  |  |  |
| Obese | 222 (18.6) |  |  |  |
| **BSA(m<sup>2</sup>)** | | Mean BSA ±<br>SD 1.88 ±<br>0.24m<sup>2</sup> | | |
| 1.20-1.50 | 55 (4.6) |  |  |  |
| 1.50-1.80 | 378 (31.7) |  |  |  |
| 1.80-2.10 | 519 (43.5) |  |  |  |
| 2.10-2.40 | 240 (20.1) |  |  |  |

### 4.2. Established baseline normal echocardiographic reference values of the left ventricular dimensions in the study population

The mean septum in diastole and systole were 8.12 ± 1.33 mm and 9.03 ± 1.37 mm respectively. The LV diameter in diastole was 44.78 ± 5.48 mm and the LV diameter in systole was 33.47 ± 5.12 mm. Post wall in diastole and systole, were 8.31 ± 1.24 mm and 9.47 ± 1.26 mm respectively. While relative wall thickness was 0.38 ± 0.08 mm, (table 4).

**Table 4:** Baseline normal echocardiographic values of the left ventricular parameters. (n = 1192).

| | Range | Median | Mean ±SD | Interquartile Range |
| --- | --- | --- | --- | --- |
| **Normal Subjects (n = 1192)** |  |  |  |  |
| Septum Diastole | 6.0-17.0 | 8.0 | $8.12 \pm 1.33$ | 7.0-9.0 |
| LV Diastole | 26.0-67.4 | 46.0 | $44.78 \pm 5.48$ | 41.0-49.0 |
| Post Wall Diastole | 6.0-14.0 | 8.0 | $8.31 \pm 1.24$ | 7.2-9.0 |
| Septum Systole | 5.0-14.3 | 9.0 | $9.03 \pm 1.37$ | 8.0-10.0 |
| LV Systole | 21.0-57.6 | 34.0 | $33.47 \pm 5.12$ | 29.5-37.0 |
| Post Wall Systole | 6.0-17.8 | 9.5 | $9.47 \pm 1.26$ | 8.9-10.1 |
| RW | 0.20-0.79 | 0.36 | $0.38 \pm 0.08$ | 0.32-0.42 |

**Table 5**. Shows normal echocardiographic values of the left ventricular diamensions according to age.

**Table 5:** Baseline normal echocardiographic values of the left ventricular parameters according to Age (n = 1192).

| Age (yrs) | Septum Diastole | Septum Systole | Post wall Diastole | Post wall Systole | LV Diastole | LV Systole | LV Mass | RWT |
| --- | --- | --- | --- | --- | --- | --- | --- | --- |
| 18-22 | 7.53±0.99* | 8.61±1.45 | 7.72±0.94* | 8.97±1.10* | 44.68±4.52 | 33.19±4.45 | 105.92±22.07* | 0.35±0.06* |
| 23-27 | 7.55±1.27 | 8.46±1.48* | 7.99±1.00 | 9.29±0.93 | 45.18±4.98 | 34.41±4.46 | 110.79±25.60 | 0.36±0.06 |
| 28-32 | 7.69±1.07 | 8.72±1.34 | 8.25±1.22 | 9.49±0.91 | 45.58±4.47 | 33.80±4.04 | 116.00±24.75 | 0.37±0.08 |
| 33-37 | 7.92±1.10 | 8.54±1.11 | 8.10±1.11 | 9.55±1.04 | 45.63±5.81 | 34.47±5.32 | 117.38±29.84 | 0.36±0.08 |
| 38-42 | 8.06±1.19 | 9.02±1.32 | 8.27±1.00 | 9.70±1.34 | 44.13±5.94 | 33.83±4.96 | 113.43±25.09 | 0.38±0.08 |
| 43-47 | 8.67±1.12<sup>+</sup> | 9.75±1.30<sup>+</sup> | 8.67±1.30<sup>+</sup> | 10.00±1.14<sup>+</sup> | 45.01±4.85 | 33.56±5.00 | 127.33±28.82<sup>+</sup> | 0.39±0.08 |
| 48-52 | 8.21±0.93 | 9.52±1.04 | 8.45±0.95 | 9.65±1.15 | 44.42±4.78 | 32.13±4.45 | 117.81±23.84 | 0.39±0.06 |
| 53-57 | 8.35±1.30 | 9.22±1.32 | 8.45±1.35 | 9.48±1.25 | 44.56±5.38 | 33.44±5.16 | 120.88±33.73 | 0.38±0.07 |
| 58-62 | 8.40±1.38 | 9.10±1.43 | 8.47±1.39 | 9.58±1.30 | 44.27±5.83 | 33.58±5.07 | 120.42±34.45 | 0.39±0.08 |
| 63-67 | 8.50±1.37 | 9.23±1.33 | 8.51±1.13 | 9.58±1.41 | 43.31±6.65* | 31.73±6.84* | 117.58±35.66 | 0.40±0.08<sup>+</sup> |
| 68-72 | 8.22±1.56 | 8.92±1.51 | 8.49±1.42 | 9.39±1.36 | 43.68±5.86 | 32.72±4.61 | 115.58±30.46 | 0.40±0.09<sup>+</sup> |
| 73-77 | 7.95±1.40 | 9.04±1.41 | 8.20±1.21 | 9.29±1.28 | 45.80±5.30 | 33.79±5.42 | 119.17±28.91 | 0.36±0.08 |
| 78-82 | 7.76±1.21 | 8.74±1.35 | 7.87±1.07 | 8.99±1.32 | 46.00±4.48 | 34.85±4.84<sup>+</sup> | 114.74±23.98 | 0.35±0.07* |
| 83-87 | 7.94±1.64 | 8.90±1.19 | 8.09±1.42 | 9.26±1.34 | 47.08±3.88<sup>+</sup> | 34.61±4.79 | 124.48±41.98 | 0.35±0.07* |
| 88-93 | 7.80±1.16 | 8.96±1.22 | 8.25±1.21 | 9.44±1.28 | 45.66±5.87 | 34.60±4.79 | 118.80±33.03 | 0.37±0.07 |
| *a = not considered for least or highest ( as n = 1); \* = least mean; + = highest mean* |  |  |  |  |  |  |  |  |

Those aged 18-22 years had the least mean of septum in diastole was 7.53 ± 0.99 mm, post wall in diastole 7.72 ± 0.94 mm, post wall systole 8.97 ± 1.10 mm and LV mass 105.92 ± 22.07kg. Those aged 43-47 years had highest mean of septum in diastole 8.67 ± 1.12 mm, septum in systole 9.75 ± 1.30 mm, post wall in diastole 8.67 ± 1.30 mm, and post wall in systole 10.00 ± 1.14 mm. Those aged 78-82 years were associated with highest mean LV diameter in systole 34.85 ± 4.84 mm, while those aged 83-87 years were associated with highest mean LV diameter in diastole 47.08 ± 3.88 mm. Highest mean in RWT was 0.40 ± 0.08 mm was among those aged 63-67 and 68-72 years (table 5).

Table 6 Baseline normal echocardiographic values of the left ventricular parameters according to Sex (n = 1,192).

**Table 6:** Baseline normal echocardiographic values of the left ventricular parameters according to Sex (n =1192).

| **M±SD** | Male | Female |
| --- | --- | --- |
| Septum Diastole | 8.05±1.27 | 8.20±1.38<sup>+</sup> |
| LV Diastole | 44.80±5.71<sup>+</sup> | 42.75±5.21 |
| Post wall Diastole | 8.32±1.26<sup>+</sup> | 7.51±1.22 |
| Septum Systole | 8.99±1.33 | 9.08±1.42<sup>+</sup> |
| LV Systole | 33.54±5.37<sup>+</sup> | 30.38±4.81 |
| Post wall Systole | 9.45±1.26 | 9.50±1.26<sup>+</sup> |
| LV Mass | 117.52±31.49 | 118.23±30.02<sup>+</sup> |
| RWT | 0.38±0.08 | 0.38±0.08 |

<sup>+</sup> = *highest mean*.

Females had higher mean septum in diastole 8.20 ± 1.38 mm, septum in systole 9.08 ± 1.42 mm, post wall in systole 9.50 ± 1.26 mm and LV mass 118.23 ± 30.02kg. Males had higher mean LV diameter in diastole 44.80 ± 5.71 mm, post wall in diastole 8.32 ± 1.26 mm and LV diameter in systole 33.54 ± 5.37 mm. Both male and femal had the RWT 0.38 ± 0.08 mm.

**Table 7**. shows the left ventricular echocardiography measurements according to BMI. The underweight had the least mean septum diastole 7.74 ± 0.84kg, LV diameter in diastole 39.80 ± 6.31mm, LV diameter in systole 29.10 ± 4.58kg, post wall in systole 9.25 ± 1.18kg and LV mass 94.31 ± 28.29kg. Highest mean septum in diastole 8.35 ± 1.33kg, post wall in systole 9.64 ± 1.36kg and LV mass 120.92 ± 31.91kg were observed among the overweight. Highest mean post wall in diastole 8.47 ± 1.34kg and septum in systole 9.26 ± 1.41kg were observed among the obese.

**Table 7:** Baseline normal echocardiographic values of the left ventricular parameters according to BMI of the participants (n = 1192).

| BMI<br>(kg/m<sup>2</sup>) | Septum<br>Diastole | LV<br>Diastole | Post wall<br>Diastole | Septum<br>Systole | LV<br>Systole | Post wall<br>Systole | LV<br>Mass | RWT |
| --- | --- | --- | --- | --- | --- | --- | --- | --- |
| Underweig<br>ht | $7.74 \pm 0.8$<br>4* | $39.80 \pm 6.$<br>31* | $8.31 \pm 1.0$<br>5 | $8.98 \pm 1.4$<br>2 | $29.10 \pm 4.5$<br>8* | $9.25 \pm 1.1$<br>8* | $94.31 \pm 28.2$<br>9* | $0.43 \pm 0.0$<br>9<sup>+</sup> |
| Normal<br>weight | $7.86 \pm 1.2$<br>8 | $45.34 \pm 5.$<br>04<sup>+</sup> | $8.15 \pm 1.1$<br>9* | $8.83 \pm 1.3$<br>2* | $33.84 \pm 4.7$<br>2<sup>+</sup> | $9.36 \pm 1.1$<br>7 | $116.02 \pm 28.$<br>88 | $0.37 \pm 0.0$<br>8* |
| Overweight | $8.35 \pm 1.3$<br>3<sup>+</sup> | $44.66 \pm 5.$<br>60 | $8.43 \pm 1.2$<br>4 | $9.16 \pm 1.3$<br>8 | $33.03 \pm 5.0$<br>1 | $9.64 \pm 1.3$<br>6<sup>+</sup> | $120.92 \pm 31.$<br>91<sup>+</sup> | $0.38 \pm 0.0$<br>7 |
| Obese | $8.33 \pm 1.3$<br>4 | $43.89 \pm 6.$<br>01 | $8.47 \pm 1.3$<br>4<sup>+</sup> | $9.26 \pm 1.4$<br>1<sup>+</sup> | $33.63 \pm 6.0$<br>3 | $9.42 \pm 1.2$<br>5 | $117.47 \pm 32.$<br>49 | $0.39 \pm 0.0$<br>9 |
| *a = not considered for least or highest ( as n = 1); \* = least mean; + = highest mean* |  |  |  |  |  |  |  |  |

**Table 8**. shows the left ventricular echocardiography measurements according to BSA.

**Table 8:** Baseline normal echocardiographic values of the left ventricular parameters according to BSA, n = 1192.

| BSA (m<sup>2</sup>) | Septum Diastole | LV Diastole | Post wall Diastole | Septum Systole | LV Systole | Post wall Systole | LVM | RWT |
| --- | --- | --- | --- | --- | --- | --- | --- | --- |
| **Normal Subjects (n = 1192)** |  |  |  |  |  |  |  |  |
| 1.20-1.50 | 8.41±1.55<sup>+</sup> | 45.17±4.59 | 7.85±1.2<br>2* | 9.67±1.27<sup>+</sup> | 32.03±4.82 | 9.33±0.9<br>8 | 118.44±42.0<br>3 | 0.35±0.07<br>* |
| 1.50-1.80 | 7.98±1.44* | 45.37±5.04<br>+ | 8.30±1.3<br>7 | 8.75±1.42* | 34.74±4.74<br>+ | 9.26±1.2<br>6* | 119.02±30.8<br>7 | 0.37±0.08 |
| 1.80-2.10 | 8.15±1.27 | 44.97±5.50 | 8.47±1.2<br>0<sup>+</sup> | 9.00±1.32 | 33.56±5.15 | 9.61±1.2<br>5<sup>+</sup> | 120.39±30.0<br>5<sup>+</sup> | 0.38±0.08<br>+ |
| 2.10-2.40 | 8.22±1.18 | 43.32±6.04<br>* | 8.10±1.0<br>6 | 9.39±1.31 | 31.60±5.07<br>* | 9.52±1.3<br>1 | 110.40±28.2<br>4* | 0.38±0.08<br>+ |
| * = least mean; + = highest mean |  |  |  |  |  |  |  |  |

BSA of 1.50-1.80 m^2^ had the least mean septum in diastole 7.98 ± 1.44 mm, septum in systole 8.75 ± 1.42 mm and post wall in systole 9.26 ± 1.26 mm. Highest mean septum diastole was 8.41 ± 1.55 mm and septum in systole was 9.67 ± 1.27 mm were observed among those with BSA of 1.20-1.50 m^2^. Those with BSA of 1.80-2.10 m^2^ had the highest mean in LV mass of 120.39 ± 30.05kg.

**Table 9**. presents the result of the relationship between age, height, weight, BMI, BSA, sex and left ventricular parameters. There was significant relationship between the following parameters. Age, had a weak negative relationship with post wall systole (r_s_ = -.066, p = .023). For height, it had a weak negative relationship with LV systole (r_s_ = -.156, p < .001) and also a weak but positive relationship with post wall systole (r_s_ = .076, p = .009). Weight had a weak positive relationship with septum diastole (r_s_ = .130, p < .001), post wall diastole (r_s_ = .058, p = .044), septum systole (r_s_ = .122, p < .001), post wall systole (r_s_ = .090, p = .002) and RWT (r_s_ = .113, p < .001), and a weak negative relationship with LV diastole (r_s_ = - .117, p < .001) and LV systole (r_s_ = -.149, p < .001). BMI had a weak positive relationship with septum diastole (r_s_ = .182, p < .001), post wall diastole (r_s_ = .124, p < .001), septum systole (r_s_ = .120, p < .001), post wall systole (r_s_ = .077, p = .008) and RWT (r_s_ = .168, p < .001) and a weak negative relationship with LV diastole (r_s_ = -.120, p < .001) and LV systole (r_s_ = -.061, p = .037). For BSA, there was weak positive relationship with septum diastole (r_s_ = .107, p < .001), septum systole (r_s_ = .109, p < .001), post wall systole (r_s_ = .098, p = .001) and RWT (r_s_ = .093, p = .001) and weak negative relationship with LV diastole (r_s_ = -.108, p < .001) and LV systole (r_s_ = -.166, p < .001). For sex, it had no significant relationship with the parameters (r_s_ = -.006, p < .001).. No other significant relationship existed.

**Table 9:** Correlation of left ventricular Parameters and Body Parameters of the participants.

|  | Age | Height | Weight | BMI | BSA | Sex |
| --- | --- | --- | --- | --- | --- | --- |
| Statistic | $r_s$ | $r_s$ | $r_s$ | $r_s$ | $r_s$ | $r_{pb}$ |
| Septum Diastole | 0.005 | -0.014 | .130* | .182* | .107* | 0.057 |
|  | **0.872** | **0.624** | **< .001** | **< .001** | **< .001** | **0.051** |
| LV Diastole | 0.051 | -0.047 | -.117* | -.120* | -.108* | -0.004 |
|  | **0.081** | **0.103** | **< .001** | **< .001** | **< .001** | **0.889** |
| Post wall Diastole | -0.001 | -0.006 | .058* | .124* | 0.041 | -0.006 |
|  | **0.972** | **0.827** | **0.044** | **< .001** | **0.156** | **0.845** |
| Septum Systole | 0.017 | 0.009 | .122* | .120* | .109* | 0.033 |
|  | **0.556** | **0.747** | **< .001** | **< .001** | **< .001** | **0.266** |
| LV Systole | 0.003 | -.156* | -.149* | -.061* | -.166* | -0.016 |
|  | **0.924** | **< .001** | **< .001** | **0.037** | **< .001** | **0.59** |
| Post wall Systole | -.066* | .076* | .090* | .077* | .098* | 0.02 |
|  | **0.023** | **0.009** | **0.002** | **0.008** | **0.001** | **0.489** |
| LVM | 0.04 | -0.051 | -0.022 | 0.029 | -0.031 | 0.012 |
|  | **0.172** | **0.081** | **0.453** | **0.309** | **0.287** | **0.691** |
| RWT | -0.013 | 0.019 | .113* | .168* | .093* | -0.006 |
|  | **0.655** | **0.511** | **< .001** | **< .001** | **0.001** | **0.849** |
| Statistics: $r_s$ = Spearman Correlation; $r_{pb}$ = Point-biserial Correlation | | | | | | |
| Correlation Coefficient (lighted); p-value (bolded); * implies significant relationship ~ significance exists at $p < .05$ | | | | | | |

Table 10, showed indexed baseline echocardiographic normal reference values of the left ventricular parameters of the participants. The mean for the indexed left ventricular parameters were distributed thus: indexed septum in diastole 4.38 ± 0.91mm, indexed septum in systole 4.86 ± 0.95mm, indexed LV diameter in diastole 24.21 ± 4.46mm, indexed LV diameter in systole 18.11 ± 3.79mm indexed post wall in diastole 4.49 ± 0.89mm, indexed post wall in systole 5.10 ± 0.90mm, and indexed LV mass 63.73 ± 19.31mm. Specifically for sex-based distribution, females had higher mean indexed septum in diastole 4.42 ± 0.9mm, indexed septum in systole 4.88 ± 0.9mm, indexed post wall in systole 5.11 ± 0.8 mm and indexed LV mass 63.94 ± 19.6mm. Males had higher indexed post wall in diastole (4.49 ± 0.90 mm), indexed LV diameter in diastole (24.22 ± 4.53mm) and indexed LV diameter in systole (18.16 ± 3.90mm).

**Table 10:**
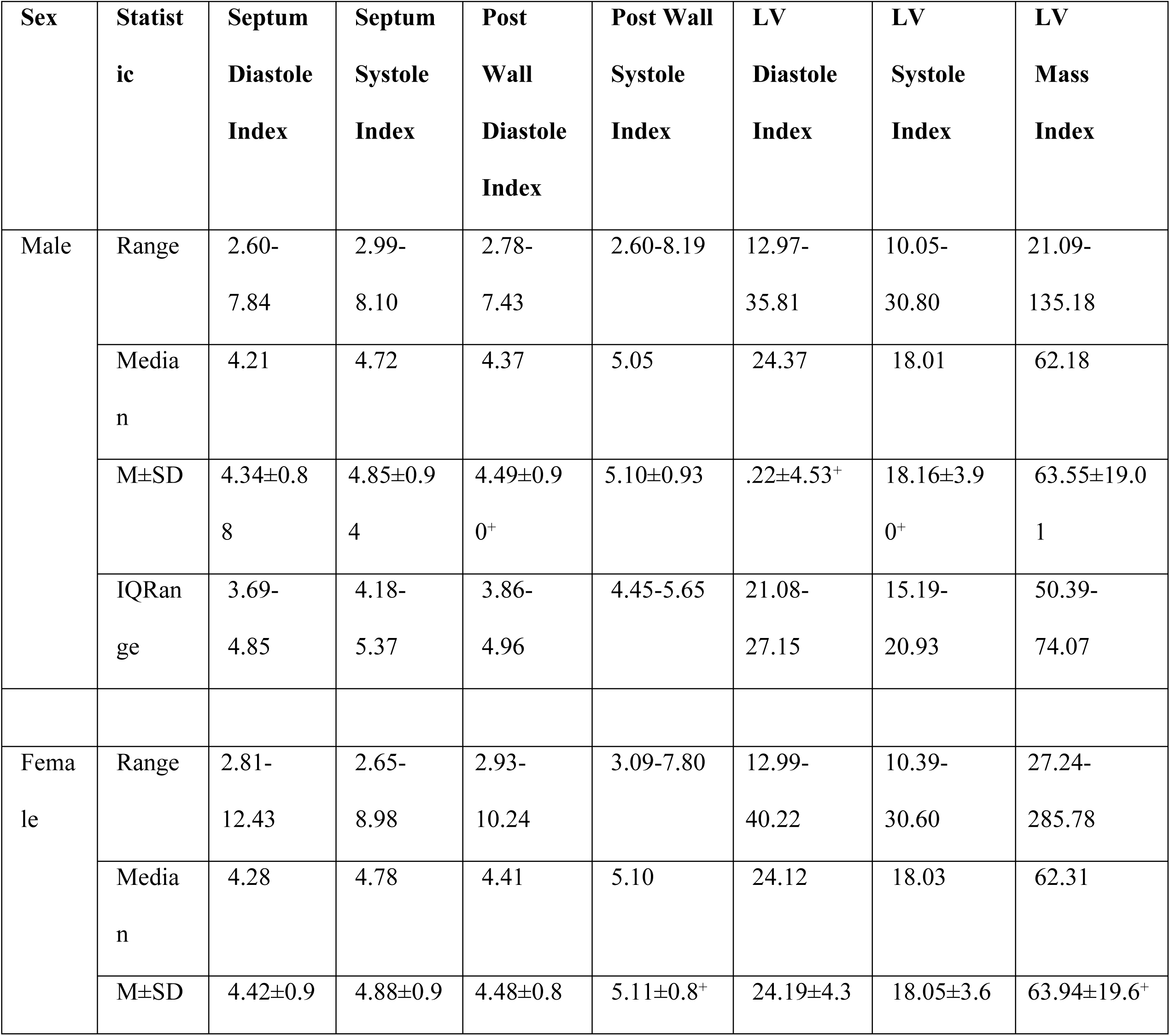
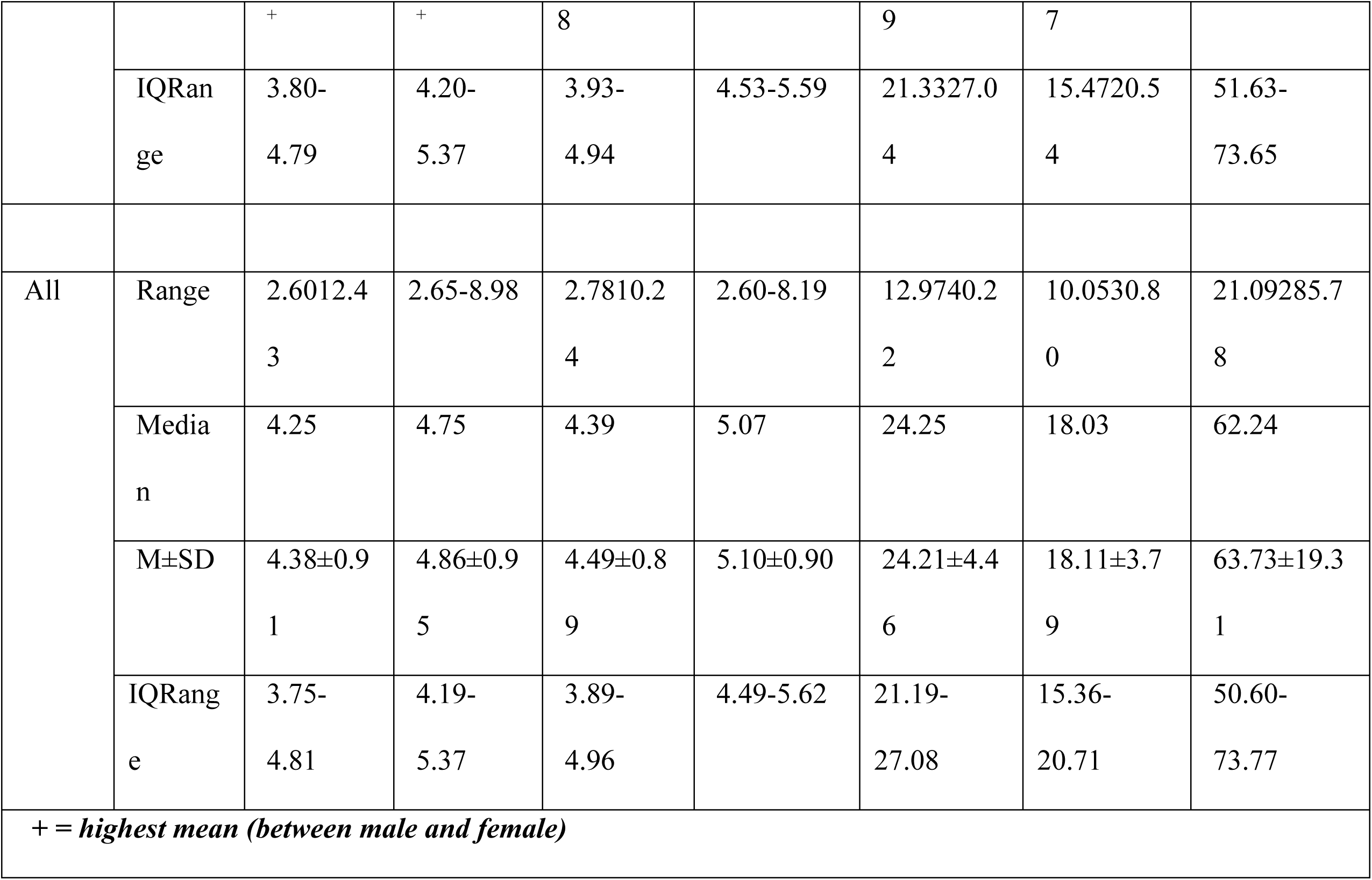
Indexed echocardiographic baseline normal values of the left ventricular parameters of the participants.

**Table 11**. Presents a comparison between Left ventricular measurements from our study and those of American society of echocardiography (ASE) and European association of cardiovascular imaging (EACVI). For LV diastole, 70.6% males and 89.7% females in our study; 79.4% subjects altogether were within the specified ASE and EACVI reference values. The indexed LV diastole was 57.0% males, 54.4% females and 55.8% subjects altogether. 89.7% males, 62.6% females and 77.1% subjects’ altogether were found within the LV systole ASE and EACVI reference values while 64.9% males, 70.3% females and 67.4% subjects altogether were found within indexed LV in systole reference values. For LV mass, the distribution was thus: 80.1% males, 92.2% females and 85.9% subjects altogether; while for indexed LV mass, it was 75.6% males, 84.8% females and 79.9% subjects. For septum diastole, 94.8% males, 82.3% females and 89.0% subjects altogether were within the specified ASE & EACVI reference values for septum diastole, while for posterior wall diastole, 93.3% males, 81.0% females and 87.6% subjects altogether were within its specified reference values.

**Table 11:** Findings of Echocardiography Measurements in Comparison to ASE and EACVI measurements.

| Parameter | Male<br>(n = 639) |  |  | Female<br>(n = 553) |  |  | Overall<br>(n = 1192) |
| --- | --- | --- | --- | --- | --- | --- | --- |
|  | ASE & EACVI | This Work | With ASE & EACVI | ASE & EACVI | This Work | Within ASE & EACVI | Within ASE & EACVI |
|  |  | M±SD<br>(Range) | n (%) |  | M±SD<br>(Range) | n (%) | n (%) |
| LV diastole | 42-58 | 44.80±5.71 | 451 | 38-52 | 44.75±5.21 | 496 | 947 |
|  |  | (27.1-67.4) | -70.6 |  | (26-64) | -89.7 | -79.4 |
| Indexed LV diastole | 22-30 | 24.22±4.53 | 364 | 23-31 | 24.19±4.39 | 301 | 665 |
|  |  | (12.97-35.81) | -57 |  | (12.99-40.22) | -54.4 | -55.8 |
| LV systole | 25-40 | 33.54±5.37 | 573 | 22-35 | 33.38±4.81 | 346 | 919 |
|  |  | (21.0-57.6) | -89.7 |  | (22-51) | -62.6 | -77.1 |
| Indexed LV systole | 13-21 | 18.16±3.90 | 415 | 13-21 | 18.05±3.67 | 389 | 804 |
|  |  | (10.05-30.80) | -64.9 |  | (10.39-30.60) | -70.3 | -67.4 |
| LV mass | 88-224 | 117.52±31.49 | 512 | 67-162 | 118.23±30.02 | 510 | 1022 |
|  |  | (45.9-234.0) | -80.1 |  | (54.5-390.8) | -92.2 | -85.9 |
| Indexed LV mass | 49-115 | 63.55±19.01 | 483 | 43-95 | 63.94±19.66 | 469 | 952 |
|  |  | (21.1-135.2) | -75.6 |  | (27.2-285.8) | -84.8 | -79.9 |
| Septum diastole | 06-Oct | 8.05±1.27 | 606 | 06-Sep | 8.20±1.38 | 455 | 1061 |
|  |  | (6-13) | -94.8 |  | (6-17) | -82.3 | -89 |
| Indexed Septum diastole | - | 4.34±0.88 | - | - | 4.42±0.94 | - | - |
|  |  | (2.6-7.8) |  |  | (2.8-12.4) |  |  |
| Septum systole | - | 8.99±1.33 | - | - | 9.08±1.42 | - | - |
|  |  | (6.0-13.0) |  |  | (5.0-14.3) |  |  |
| Indexed Septum systole | - | 4.85±0.94 | - | - | 4.88±0.96 | - | - |
|  |  | (3.0-8.1) |  |  | (2.7-9.0) |  |  |
| Post wall diastole | 06-Oct | 8.32±1.26 | 596 | 06-Sep | 8.31±1.22 | 448 | 1044 |
|  |  | (6-13) | -93.3 |  | (6-14) | -81 | -87.6 |
| Indexed Post wall diastole | - | 4.49±0.90 | - | - | 4.48±0.88 | - | - |
|  |  | (2.8-7.4) |  |  | (2.9-10.2) |  |  |
| Post wall systole | - | 9.45±1.26 | - | - | 9.50±1.26 | - | - |
|  |  | (6.0-17.8) |  |  | (6.0-17.0) |  |  |
| Indexed Post wall systole | - | 5.10±0.93 | - | - | 5.11±0.87 | - | - |
|  |  | (2.6-8.2) |  |  | (3.1-7.8) |  |  |

## 5. Discussion

### 5.1 Baseline normal Echocardiographic Values of the Left Ventricle

In this study, various measurements of the left ventricular dimensions in apparently normal subjects were obtained and indexed to BSA. This agrees with findings from other similar studies which reported that indexing echocardiographic parameters to BSA produces more reliable data [2, 6]. The study by [10] described only the profile of cardiomyopathy in Nigeria among children in Jos Northern part of Nigeria. Another study by [9] in western Nigeria also described only the profile of cardiomyopathy in Nigeria among children in the region. Thus there is still need for echocardiographic normal values in adult population in Nigeria.

In this study, 1192 apparently healthy subjects were studied to establish baseline normal reference values of left ventricular dimensions. In general it was discovered that left ventricular dimensions increases with increase in age and morphology until mid age in this present study. This was in agreement with the work done by [2] which stated that the size and morphology of the left ventricle varies with age and from person to person. This study, ensured inclusion of nearly 97% of subjects (from the 3rd to 97^th^ percentile) in producing the baseline normal echocardiographic values of the study population. This implied that there was adequate subject representation in the study. The mean and standard deviation of left ventricular internal diameter in diastole (LVEDD) and systole (LVESD) were 44.78 ± 5.48 mm and 33.47 ± 5.12 mm, respectively. males had larger LVEDD (44.80±5.71 mm), LVESD (33.54±5.37 mm), and PWd (8.32±1.26 mm) than females (p < 0.001). Left ventricular mass (118.23 ± 30.02 kg), IVSd (8.20 ± 1.38 mm) and IVSs (9.08 ± 1.42 mm) were higher in females (p < 0.001). These were similar with a study on normal reference values of echocardiographic measurements in young Egyptian adults where males also had larger values than females in these parameters [2].

### 5.2 Correlation of age, height, weight, BMI and BSA, with left ventricular echocardiographic parameters

Age had a weak negative relationship with left ventricular posterior wall in systole (r_s_ = - 0.066, p = 0.023) while height, had a weak negative relationship with LV end -systolic diameter (r_s_ = - 0.156, p < 0.001) Weight had a weak positive relationship with interventricular septum in diastole (r_s_ = .130, p < .001),Relative wall thickness has a weak negative relationship with LV end-diastolic diameter (r_s_ = -.117, p < .001). These implied that these measurements can been taken at any age, height and weight. Body mass index had a weak positive relationship with interventricular septum in diastole (r_s_ = .182, p < .001), left ventricular posterior wall in diastole (r_s_ = .124, p < .001), interventricular septum in systole (r_s_ = .120, p < .001), left ventricular posterior wall in systole (r_s_ = .077, p = .008) and RWT (r_s_ = .168, p < .001). However, BMI had a weak negative relationship with LV end-diastolic diameter (r_s_ = -.120, p < .001) and LV end -systolic diameter (r_s_ = -.061, p = .037).

Body surface area had weak positive relationship with interventricular septum in diastole (r_s_= .107, p < .001), interventricular septum in systole (r_s_ = .109, p < .001), left ventricular posterior wall in systole (r_s_ = .098, p = .001) and RWT (r_s_ = .093, p = .001). BSA had weak negative relationship with LV end-diastolic diameter (r_s_ = -.108, p < .001) and LV end - systolic diameter (r_s_ = -.166, p < .001). The study showed that a linear correlation existed between all echocardiographic measurements with age, height, weight, BMI and BSA which is similar to the findings of [11]. These imply that echocardiographic parameters can be indexed to body surface area to obtain a more reliable normal reference values in our locality.

### 5.3 Baseline normal echocardiographic values indexed to BSA

Producing normal values for dimensions and functions of the heart is very important to avoid misclassification of normal persons into the high risk category and the reverse [12] Studies have shown that using measurements indexed to BSA provides more reliable information [13]. In this study, participants’ absolute LV dimesions and LV mass were smaller than the ASE-recommended normal values. These differences were significantly minimized when the values were indexed to BSA. These findings imply that the practice of using absolute values for defining normality of various cardiac chamber dimensions in our locality should be discouraged. The BSA-indexed left ventricular baseline normal reference values in our locality should be used during echocardiography investigations because there is well-developed fact that indexing allows comparisons in subjects with different body sizes [14, 15, 8].These changes have also been observed among different races and ethnicities which raised the importance of this study keeping in mind that today’s baseline normal values are widely based on data from North American cohorts obtained in the 1970s and 80s [16, 17, 18, 4].

Specifically, males had higher indexed post wall diastole, indexed LV diameter in diastole (LVEDD) and indexed LV diameter in systole (LVEDS). Females had higher mean in indexed septum diastole, indexed septum systole, indexed post wall systole (5.11 ± 0.8) and indexed LV mass (63.94 ± 19.6). Therefore males had larger left ventricular dimensions than females in this study.

### 5.4 Comparing the study values to international reference values

Measurements obtained from this study were compared with the reference values of American Society of Echocardiography (ASE) and the European Association of Cardiovascular Imaging (EACVI). Thus for LVEDD, 70.6% were males and 89.7% were females, while 79.4% subjects altogether were within the specified ASE and EACVI reference ranges. The indexed LV in diastole was thus: 57.0% males, 54.4% females and 55.8% subjects altogether. Moreover, 89.7% males, 62.6% females and 77.1% subjects’ altogether were found within the LV systole ASE and EACVI reference range while 64.9% males, 70.3% females and 67.4% subjects altogether were found within indexed LV systole reference values.

For LV mass, the distribution was thus: 80.1% males, 92.2% females and 85.9% subjects altogether; while for indexed LV mass, it was thus: 75.6% males, 84.8% females and 79.9% subjects. For septum diastole, 94.8% males, 82.3% females and 89.0% subjects altogether were within the specified ASE and EACVI reference ranges, while for post wall diastole, 93.3% males, 81.0% females and 87.6% subjects altogether were within the specified reference ranges.

On examining these results, it was found that our locality LV dimensions were smaller than those of the caucasian population. The males in this study had a higher upper reference limit of normal for LVEDD, LVESD and IVSd. Females had a higher upper reference limit of normal for indexed LV mass. These confirmed that in this study, males had higher indexed left ventricular internal diameters while the females had thicker left ventricular walls.

## 6. Limitations of the study

Delineation of the endocardial margins on the M-Mode tracing is not always easy. The M-Mode line lies between the papillary muscle and the mitral valve may be mistaken for the posterolateral wall. During the study, there was the problem of inward motion of the small ventricles (wall and portions of the papillary muscle) from the sides, which enters the field of view of the M-Mode. This may cause the investigator to underestimate end-systolic diameter. Moreover, poor image quality and oblique views made it difficult to perform measurements. Also, this work did not measure LV volume which is obtained by 2D measurement of LV length and width, thus further studies should adopt 2D measurement of the LV volume as this work used M-Mode tracing.

## 7. Conclusion

This study established data for baseline normal values for normal left ventricle of the heart in a population of one thousand one hundred and ninety two (1,192) by echocardiography. The study, ensured inclusion of nearly 97% of subjects (from the 3rd to 97^th^ percentile). There was adequate subject representation in the study. In general, left ventricular dimensions increases with increase in age and morphology until mid age. The mean of left ventricular internal diameter in diastole (LVEDD) and systole (LVESD) were 44.78 ± 5.48 mm and ± 5.12 mm, respectively. All age group and gender were adequately represented with slightly more male participants.

Males had larger LVEDD (44.80±5.71 mm), LVESD (33.54±5.37 mm), and PWd (8.32±1.26 mm) than females (p < 0.001). Left ventricular mass (118.23 ± 30.02 kg), IVSd (8.20 ± 1.38 mm) and IVSs (9.08 ± 1.42 mm) were higher in females (p < 0.001).

The mean for the indexed echocardiography parameters established by the study were as follows: indexed LV diastole (24.21 ± 4.46mm) and indexed LV systole (18.11 ± 3.79mm). Indexed LV mass (63.73 ± 19.31kg), indexed septum diastole (4.38 ± 0.91mm) and indexed septum systole (4.86 ± 0.95mm). Indexed post wall diastole (4.49 ± 0.89mm), and indexed post wall systole (5.10 ± 0.90mm). Specifically, males had higher indexed post wall diastole (4.49 ± 0.90mm), indexed LV diameter in diastole (LVEDD) (22.37 ± 4.53mm) and indexed LV diameter in systole (LVEDS) (18.16 ± 3.90mm). Females had higher mean in indexed septum diastole (4.42 ± 0.9mm), indexed septum systole (4.88 ± 0.9mm), indexed post wall systole (5.11 ± 0.8mm) and indexed LV mass (63.94 ± 19.6mm). Therefore males had larger left ventricular dimensions than females in this study.

The values of LV dimensions and LV mass of this study were smaller when compared with international baseline reference values when indexed to BSA. This was attributed to racial difference. A linear correlation existed between all echocardiographic measurements with age, height, weight, BMI and BSA in the study population.

Finally, this research established baseline normal reference values of left ventricular dimensions in the locality by echocardiography.

## Acknowledgements

We will ever remain grateful to the management and staff of Waves medical diagnostic and research center Nnewi, Anambra State Nigeria for their assistance and co-operation in using their facility during the data collection. Many thank to Mr Charles Ugwu (Statistician) and his staff members for the data analysis.

## Ethics approval and consent to participate

The permission of the Ethics Committee of the Nnamdi Azikiwe University Teaching Hospital Nnewi was obtained in addition to informed consent from all the participants. Reference number: NAUTH/CS/66/vol.9/162/2016/138. Approval date on the ethical letter was on 13^th^ March 2017.

## Availability of data and material

The dataset generated and analysed during the current study are not publicly available but are available from the corresponding author on reasonable request.

## Patient consent for publication

Not applicable.

## List of abbreviations

Not applicable.

## Human rights statements and informed consent

All procedures followed were in accordance with the ethical standards of the responsible committee on human experimentation (institutional and national) and with the Helsinki Declaration of 1964 and later revisions. Informed consent was obtained from all patients for being included in the study.

## Figures

**Fig. 1:**
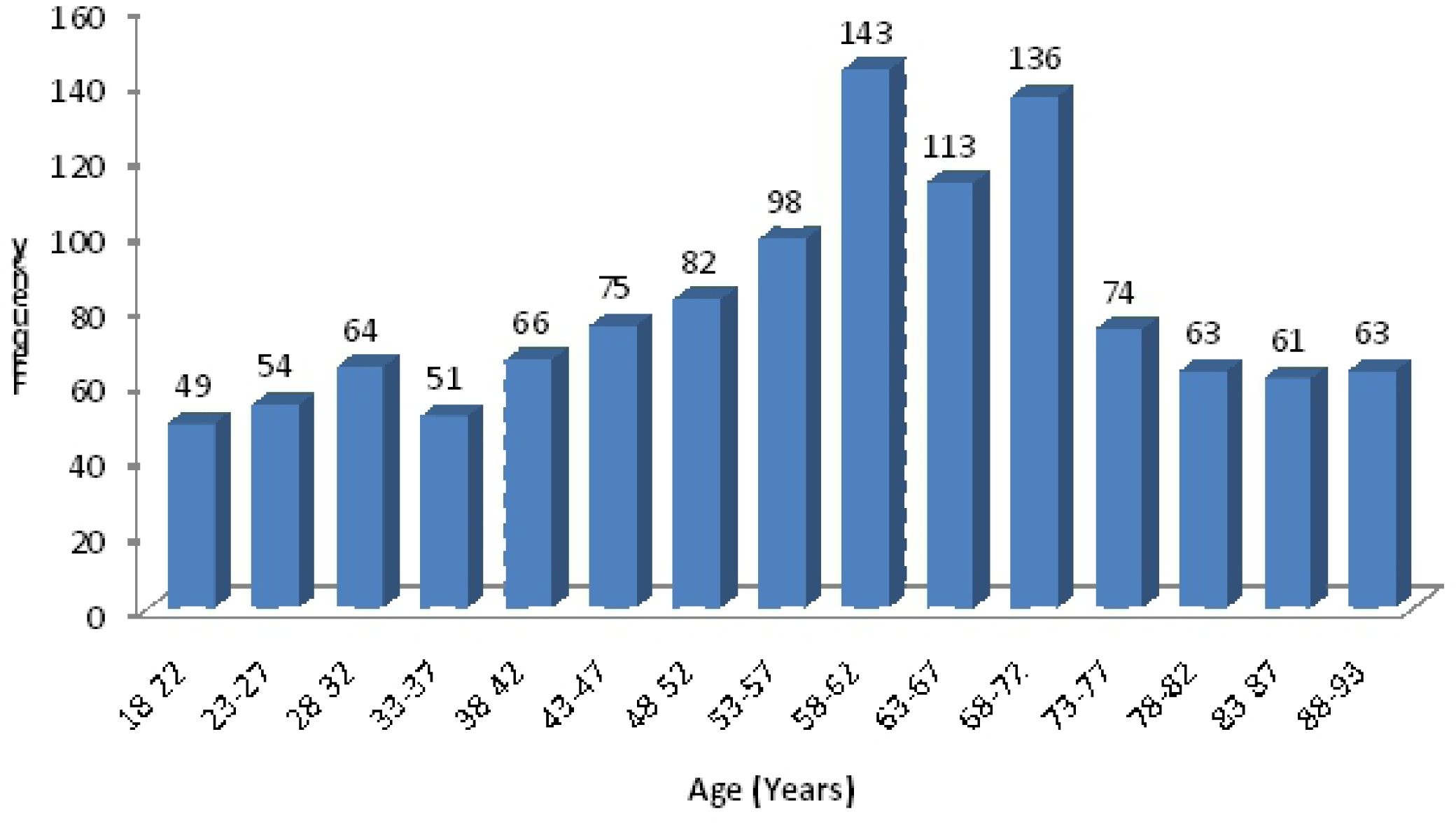
Age distribution of the participants.

**Fig. 2:**
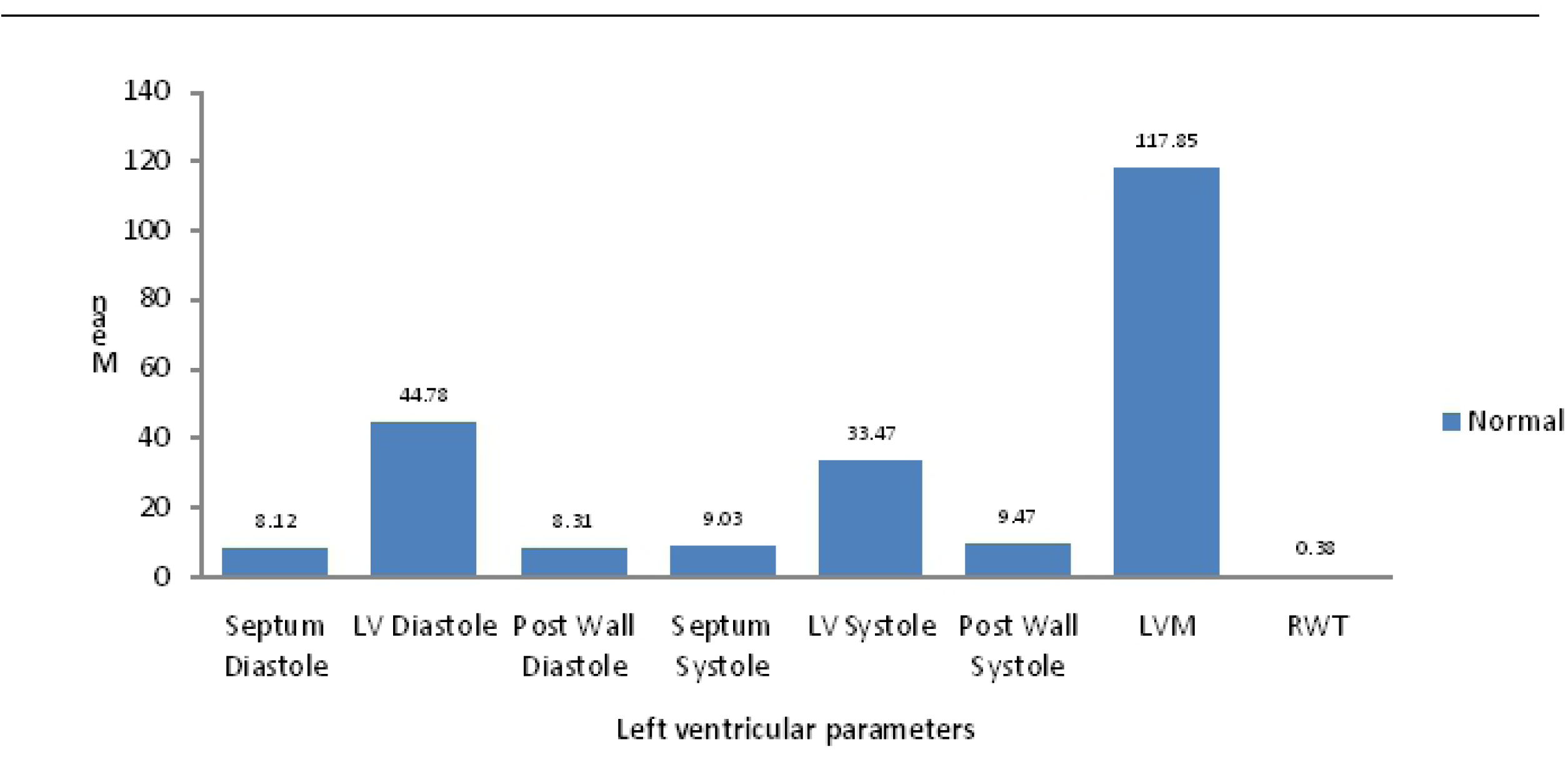
Baseline normal echocardiographic reference value of the left ventricular diamension in the study population.

## References

1. Sonography. Size of the left ventricle. Medical University of Vienna. 2017; 123 Sonography.com,

2. Ahmed Mohamed El Missiri, Khaled Abdel Lateef El Meniawy, Sherif Abdel Salam Sakr, et al.,. Normal reference values of echocardiographic measurements in young Egyptian adults. The Egyptian Heart Journal. 2016; 68: 209–215.

3. Lancellotti P, Badano LP, Lang RM, et al. Normal reference ranges for echocardiography: rationale, study design, and methodology (NORRE study). European Heart Journal of Cardiovascular Imaging, 2013; 14: 303–8.

4. Lancellotti P. Normal reference ranges for echocardiography: do we really need more?. European Heart Journal of Cardiovascular Imaging 2014; 15:253–4.

5. Hamayak, S. Cardiomyopathies: Evolution of pathogenesis concepts and potential for new therapies. World journal of cardiology. 2014; 6(6): 478–494.

6. Manish Bansal, Jagdish C. Mohan, Shantanu P. et al. Normal echocardiographic measurements in Indian adults: How different are we from the western populations? A pilot study. Indian heart journal. 2016; 68: 772–775.

7. Rudski LG, Lai WW, Afilalo J, et al. Guidelines for the echocardiographic assessment of the right heart in adults: a report from the American Society of Echocardiography endorsed by the European Association of Echocardiography, a registered branch of the European Society of Cardiology, and the Canadian Society of Echocardiography. Journal of American Society of Echocardiographers 2010; 23:685–713.

8. Lang RM, Badano LP, Mor-Avi V, et al. Recommendations for cardiac chamber quantification by echocardiography in adults: an update from theAmerican society of echocardiography and the European association of cardiovascular imaging. Journal of American Society of Echocardiography 2015; 28:1–39.

9. Animasahun, BA, Madise, AD, Ogunkunle, OO, et al. A descriptive study about dilated cardiomyopathy in children in tertiary hospital in Nigeria. Journal of clinical and experimental research in cardiology 2015; 2:1: 102.

10. Bode-Thomas F, Olukemi O, and Ige, CY. Childhood acquired heart disease Jos, Nigeria. Nigeria medical journal. 2013; 54:1:51-58.

11. Motz R, Schumacher M, Nürnberg J, et al. Echocardiographic measurements of cardiacdimensions correlate better with body length than with body weight or body surface area. Pediatric Cardiology 2014; 35:1327–36.

12. Vasan RS, Levy D, Larson MG, et al. Interpretation of echocardiographic measurements: a call for standardization. American Heart Journal 2000;139:412–22.

13. Pfaffenberger S, Bartko P, Graf A, et al. Size matters, Impact of age, sex, height, and weight on the normal heart size. Circulation Cardiovascular Imaging 2013; 6:1073–9.

14. De Simone G, Kizer JR, Chinali M, et al. Normalization for body size and population-attributable risk of left ventricular hypertrophy: the strong heart study. American Journal of Hypertension 2005;8:191–6.

15. Chirinos JA, Segers P, De Buyzere ML, et al. Left ventricular mass: allometric scaling, normative values, effect of obesity, and prognostic performance. American journal of Hypertension 2010; 56:91–8

16. Ilercil A, O’Grady MJ, Roman MJ, et al. Reference values for echocardiographic measurements in urban and rural populations of differing ethnicity: the strong heart study. Journal of American Society of Echocardiography 2001;14:6:601–11.

17. Daimon M, Watanabe H, Abe Y, et al. Normal values of echocardiographic parameters in relation to age in a healthy Japanese population: the JAMP study. Circulation Journal, 2008; 72:1859–66.

18. Chahal NS, Lim TK, Jain P, et al. Ethnicity-related differences in left ventricular function, structure and geometry: a population study of UK Indian Asian and European white subjects. Heart 2010; 96:466–71.

